# The Small GTPase Rho5 – Yet Another Player in Yeast Glucose Signaling

**DOI:** 10.1101/2024.07.29.605593

**Authors:** Franziska Schweitzer, Linnet Bischof, Stefan Walter, Silke Morris, Hans-Peter Schmitz, Jürgen J. Heinisch

## Abstract

The small GTPase Rho5 has been shown to be involved in regulating the Baker’s yeast response to stress on the cell wall, high medium osmolarity, and reactive oxygen species. These stress conditions trigger a rapid translocation of Rho5 and its dimeric GDP/GTP exchange factor (GEF) to the mitochondrial surface, which was also observed upon glucose starvation. We here show that *rho5* deletions affect carbohydrate metabolism both at the transcriptomic and the proteomic level, in addition to cell wall and mitochondrial composition. Epistasis analyses with deletion mutants in components of the three major yeast glucose signaling pathways indicate a primary role of Rho5 upstream of the Ras2 GTPase in cAMP-mediated protein kinase A signaling. We also observed an inhibitory role of Rho5 on respiratory capacity, which may be explained by its role in mitophagy.

## 1. Introduction

The yeast *Saccharomyces cerevisiae* has been employed by mankind since thousands of years for making bread and alcoholic beverages like beer and wine (Barnett, 2003). It was thus continuously selected for efficient sugar utilization, with glucose as the preferred carbon source (Borneman and Pretorius, 2015). Besides a large family of hexose transporters (Boles and Hollenberg, 1997), this prompted the evolution of complex signaling networks to detect and properly react to the sugar concentration in the medium (see Busti et al., 2010; Creamer et al., 2022; Plank, 2022; Zaman et al., 2008, for some selected reviews). As outlined in Fig. 1, three major signaling pathways have been identified in this context, which are characterized by the trimeric SNF1/AMPK complex, the Rgt2/Snf3 sensor pair, and the cAMP-activated protein kinase A (cAMP/PKA).

**Figure 1.**
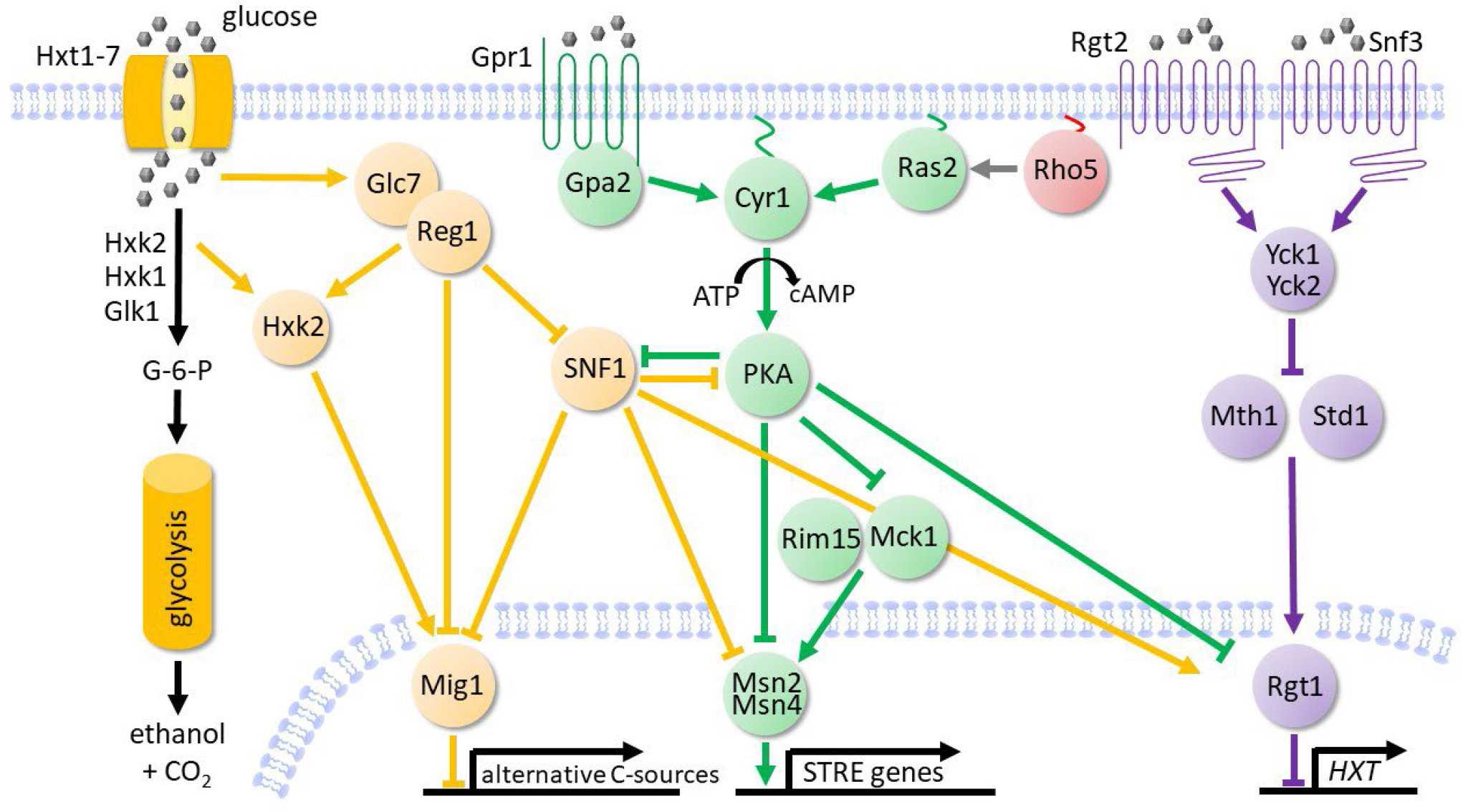
Simplified scheme of glucose signaling pathways in *Saccharomyces cerevisiae*. Glucose (grey hexagons) is internalized by hexose transporters, with Hxt1-Hxt7 contributing the major activity. After activation by hexokinase (primarily Hxk2) it is channeled into glycolysis. At high extracellular glucose concentrations the Reg1-Glc7 phosphatase complex dephosphorylates the SNF1 complex, as well as Hxk2 and Mig1, leading to repression of genes required for the utilization of alternative carbon sources (SNF1 pathway, designated in orange).

In the cAMP/PKA pathway (depicted in green) the G-protein coupled receptor Gpr1 senses extracellular glucose and transmits the signal to Gpa2, which activates the adenylate cyclase Cyr1. Cyr1 can also be activated by the redundant Ras1 and Ras2 GTPases in response to intracellular glucose-induced changes. Activated Cyr1 produces cyclic AMP which binds to the regulatory subunits of the heterotetrameric protein kinase A (PKA), triggering their dissociation from and activation of the catalytic subunits. The protein kinases Rim15 and Mck1 are inactivated by PKA-dependent phosphorylation, as are the redundant transcription factors Msn2 and Msn4. The latter control the expression of stress-response genes by binding to conserved STRE sequences in their promoters. A third glucose-responsive pathway is initiated by the Rgt2/Snf3 sensors (shown in violet), which perceive extracellular glucose and activate the yeast casein kinases Yck1 and Yck2. These phosphorylate and thereby mark the co-repressors of the transcription factor Rgt1 for proteolytic degradation, namely Mth1 and Std1. The trimeric transcription complex Rgt1/Mth1/Std1 governs the expression of several hexose transporter genes (*HXT*s). Lines ending in bars designate inhibition, arrows indicate activation of target proteins. The proposed role of Rho5 as a positive regulator of Ras2, deduced from the results of this work, is also shown.

The *SNF1* gene was originally identified in screens for yeast mutants impaired in catabolite repression for utilization of sucrose and other carbon sources than glucose (hence “**s**ucrose non-fermenters”, also designated as *CAT1*; Carlson et al., 1981; Hedbacker and Carlson, 2008; Zimmermann et al., 1977). It was then cloned and characterized as encoding the protein kinase subunit of a trimeric complex, compresing an additional gamma-subunit (Snf4), and one of three alternative beta-subunits (Gal83, Sip1, Sip2; Celenza and Carlson, 1984). Homologues of this trimeric complex in plants and animals, known as AMP-activated protein kinase (AMPK), were also found to govern energy metabolism, with malfunctions having severe effects on human health (Carling, 2017; Hardie et al., 2012). In yeast, glucose deprivation leads to phosphorylation and activation of the Snf1 kinase subunit by one of three protein kinases (Elm1, Sak1, Tos3), upon which the trimeric complex comprising Gal83 as a ß-subunit enters the nucleus and triggers gene expression for the utilization of alternative carbon sources (see Hedbacker and Carlson, 2008 and Zaman et al., 2008, and references therein, for a detailed overview). Two of its major target proteins are the transcriptional activator Adr1 and the transcriptional repressor Mig1, with the latter being exported from the nucleus upon its phosphorylation. In addition, SNF1 activity participates in many cellular interactions, which related to this work include the transcription factors Msn2/Msn4, the cAMP-activated protein kinase A (PKA), nutrient signaling through the TORC1 complex, and hexokinase PII (Coccetti et al., 2018; Lubitz et al., 2015). Interestingly, Hxk2 also functions as a co-repressor together with Mig1 in nuclear gene expression (Vega et al., 2016). Moreover, yeast cell wall synthesis was found to be regulated by the SNF1 complex in a Mig1-dependent manner (Backhaus et al., 2013; Rippert et al., 2017).

Glucose signaling mediated by the Snf3/Rgt2 sensors in *S. cerevisiae* acts on the expression of several hexose transporter genes through inactivation of a trimeric repressor complex comprising Rgt1 as the DNA-binding subunit (Fig. 1; reviewed in Busti et al., 2010 and Zaman et al., 2008). While transcription of the target *HXT* genes is repressed by limiting glucose concentrations, ample glucose triggers the proteosomal degradation of the Mth1 and Std1 subunits and thereby inactivates the repressor complex. This pathway is crosslinked to SNF1 signaling indirectly by Mig1-mediated repression of genes encoding Rgt1 and its co-repressor Mth1 (Kayikci and Nielsen, 2015), and directly by phosphorylation and activation of Rgt1 (Palomino et al., 2006).

Finally, Rgt1 repression is also alleviated upon its hyperphosphorylation by protein kinase A (PKA; Flick et al., 2003; Roy et al., 2013), the central component of the third and probably most extensively studied route of yeast glucose signaling (Fig. 1; see again Busti et al., 2010 and Zaman et al., 2008 for general overviews). In brief, glucose signaling in yeast was originally related to the action of the small GTPase homologues of human Ras (Ras1 and Ras2) in stimulating adenylate cyclase (Cannon et al., 1986; Sass et al., 1986). Much later, signaling through the G protein coupled receptor (GPCR) Gpr1 and the GTPase Gpa2 was proposed to be the more important trigger of adenylate cyclase activity (Kraakman et al., 1999). In both cases the resulting peak in cAMP concentration leads to dissociation of the inhibitory Bcy1 subunits from the tetrameric PKA complex and concomitant liberation of the catalytic subunits, with the three isoforms Tpk1, Tpk2, and Tpk3 (Toda et al., 1987a; Toda et al., 1987b). These show partially overlapping but also distinct specificities towards a variety of cytosolic target proteins and nuclear transcription factors (reviewed in Portela and Rossi, 2020; Thevelein and de Winde, 1999). Amongst the latter, the redundant transcription factors Msn2/Msn4 are a major target. They are inactivated by PKA-dependent phosphorylation and exported from the nucleus at high extracellular glucose concentrations (Boy-Marcotte et al., 1998; Durchschlag et al., 2004). Under glucose or other nutrient limitations, as well as in response to different environmental stresses (governed by the ESR pathway), they reside in the nucleus, where they activate the expression of genes through binding to stress responsive promoter elements (STREs; Schmitt and McEntee, 1996). In addition to PKA-mediated glucose signaling, nuclear export of Msn2 can also be provoked by its phosphorylation by the protein kinase Rim15, which provides a link to TORC1-mediated nutrient signaling and the regulation of autophagy (Lee et al., 2013; Yorimitsu et al., 2007).

Although this intricate glucose signaling network has been extensively studied in the model yeast *S. cerevisiae* for almost five decades, evidence for new crosstalks is constantly arising. Thus, we found that the small GTPase Rho5 may also take part in glucose and other nutrient signaling, as *rho5* mutants show strong synthetic defects with *gpa2*, *gpr1*, or *sch9* deletions (Schmitz et al., 2018). Historically, Rho5 was originally identified as a negative regulator of cell wall integrity (CWI) signaling (Schmitz et al., 2002), and found to regulate the opposing high osmolarity glycerol (HOG) pathway (Annan et al., 2008). Besides their specific functions, both pathways are also required for a proper yeast mitophagy (Mao et al., 2011). Moreover, upon exposure to oxidative stress Rho5 rapidly translocates from the plasma membrane to mitochondria, and triggers mitophagy and apoptosis (Schmitz et al., 2015; Singh et al., 2008). Interestingly, the roles of Rho5 in energy metabolism and mitochondrial functions appear to be conserved in its human homologue Rac1, whose malfunctions are associated with several diseases, including diabetes, cancer, and neurodegenerative disorders (reviewed in Bischof et al., 2024a).

In this work, both the transcriptome and proteome were analyzed in *rho5* mutants and compared to wild-type cells, which substantiated the notion that the small GTPase participates in yeast glucose signaling. We then embarked on more detailed epistasis analyses with deletion mutants in selected components of the three major signaling pathways. Results obtained are consistent with Rho5 acting primarily through the cAMP/PKA pathway.

## 2. Results

### 2.1. RNAseq and proteome analyses relate Rho5 to glucose signaling and general stress response

As Rho5 has been found to be involved in the regulation of a large number of signaling processes (Hühn et al., 2020), we decided to apply two global assays to assess the effect of *rho5* deletion mutants on yeast physiology. First, data on the transcriptome were obtained by RNAseq for the wildtype and the deletion mutant under standard growth conditions in synthetic medium with 2% glucose, both in the absence and presence of 0.8 mM hydrogen peroxide. A total of 99 genes showed a significant upregulation in their expression under standard growth conditions when the deletion mutant was compared to the wild-type, whereas 95 genes appeared to be downregulated (Fig. 2A; cutoffs applied at p-values less than 0.05 and at least a twofold change in expression; see supplementary Table S1 for a complete list of the RNAseq results).

**Figure 2.**
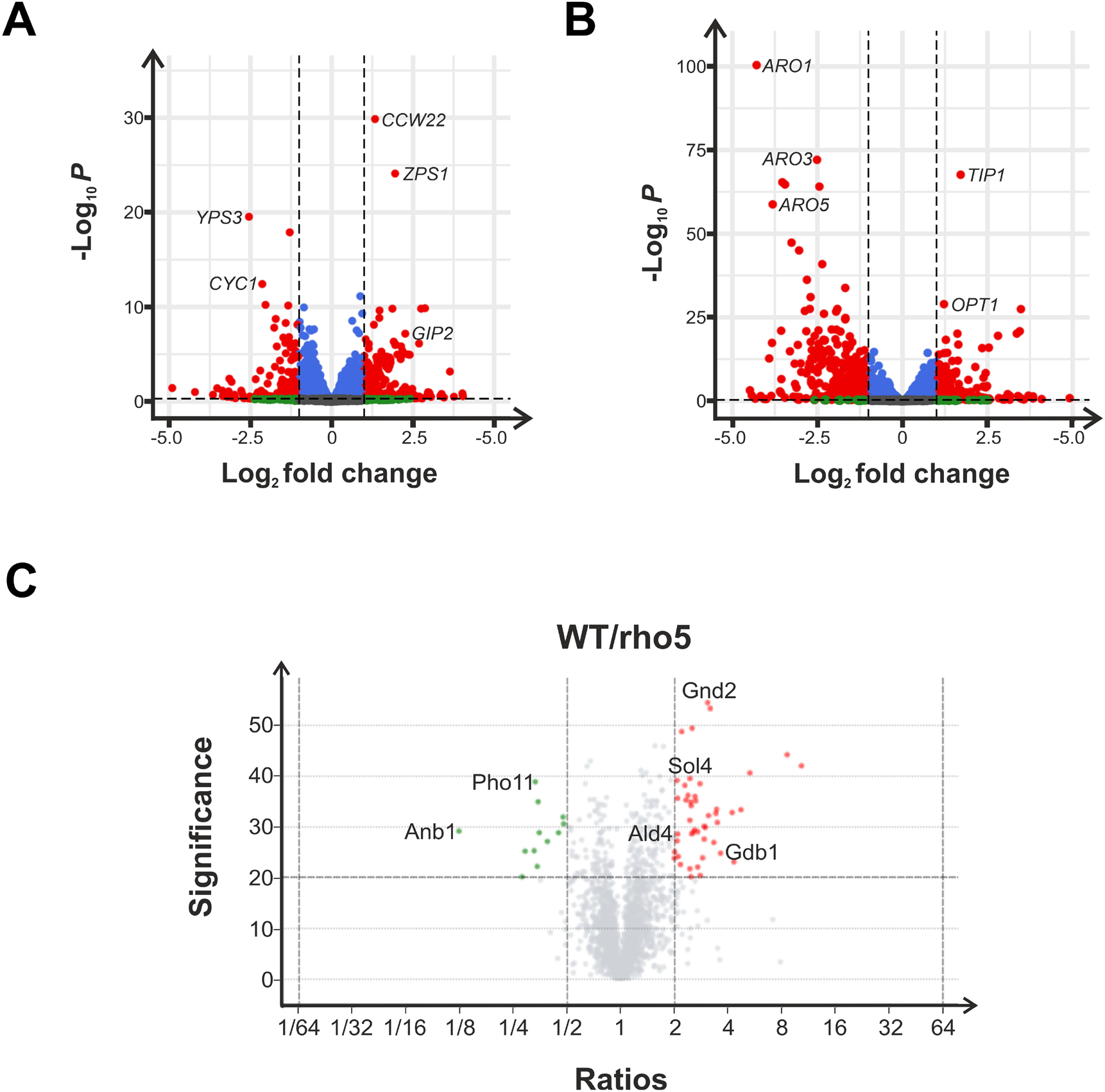
Transcriptome and proteome analyses of rho5 deletions as compared to wild-type cells. A) Volcano plot of RNAseq data comparing a rho5 deletion (FSO62-7A) to its isogenic wild-type strain (HD56-5A). Cells were grown on synthetic complete medium with 2% glucose (SCD) and general cut-offs for differentially expressed genes were applied at p-values less than 0.05 and an at least twofold change in transcript abundance, with three biological replicates for each strain. Significant expression changes are depicted in red. Transcripts not meeting the stringent cut-off criteria are designated as follows: Blue colour indicates expression changes with p-values below 0.05 but a fold-change of less than 2. Shown in green are expression changes with a fold-change of at least 2, but a p-value higher than 0.05. Grey transcripts were deemed less significant, as they have p-values higher than 0.05 and a less than twofold change. B) Volcano plot of RNA-sequencing results in which rho5 deletion cells (FSO62-7A) were compared to wild type cells (HD56-5A) after growth on SCD in the presence of 0.8 mM H2O2. Cut-offs and colour codes were applied as in A), again with three biological replicates for each strain. Volcano plot of proteins detected in mass spectrometry, comparing a rho5 deletion (FSO62-7A) to wild type cells (HD56-5A) after growth on SCD. Cut-offs were set at p-values less than 0.05 and at least a twofold change in protein abundance. Three biological replicates were used for each strain. Shown in green are all significantly down-regulated proteins, while significantly up-regulated proteins are shown in red. All proteins depicted in grey either have a p-value higher than 0.05 or a fold-change of less than 2.

As would be expected from the established Rho5 functions, the genes that were upregulated in the *rho5* deletion as compared to the wild type included those encoding cell wall and mitochondrial proteins (Table 1; note that some other genes in these two catagories were significantly down-regulated). In addition, several genes related to carbohydrate metabolism were upregulated, including several hexose transporter genes (*HXT*s) and the glucose-repressed gene *HXK1*, which encodes hexokinase I, required for sugar consumption in the late phase of wine fermentations. Moreover, genes encoding key enzymes of the pentose phosphate pathway, stress protection and reserve carbohydrate metabolism were found to be upregulated in the *rho5* deletion as compared to the wild type. Of note, many of these genes and those placed into the other metabolic groups carry stress-responsive elements (STREs) in their promoters (Table 1).

**Table 1.** Selected genes/proteins differentially expressed in a *rho5* deletion as compared to wild type after growth on synthetic complete medium with 2% glucose

| Gene name | Protein function | RNA Seq [fold change] | RNA Seq [p-value] | Mass Spec [fold change] | Mass Spec [p-value] | STRE<sup>b</sup> |
| --- | --- | --- | --- | --- | --- | --- |
| Glucose uptake and activation |  |  |  |  |  |  |
| *HXK1* | Hexokinase isoenzyme 1 | 2.07 | 1.10 × 10<sup>-2</sup> | 2.96 | 1.80 × 10<sup>-3</sup> | Yes |
| *HXT11* | Hexose transporter | 0.01 | 1.96 × 10<sup>-6</sup> | nd | nd |  |
| *HXT5* | Hexose transporter with moderate glucose affinity | 5.04 | 1.03 × 10<sup>-5</sup> | nd | nd | Yes |
| *HXT6*<sup>a</sup> | High-affinity glucose transporter | 3.87 | 1.05 × 10<sup>-6</sup> | 11.86 | 1.05 × 10<sup>-3</sup> |  |
| *HXT7*<sup>a</sup> | High-affinity glucose transporter | 6.73 | 1.53 × 10<sup>-10</sup> | 11.86 | 1.05 × 10<sup>-3</sup> |  |

| Pentose Phosphate Pathway |  |  |  |  |  |  |
| --- | --- | --- | --- | --- | --- | --- |
| *GND2* | 6-phosphogluconate dehydrogenase | 3.34 | $3.70 \times 10^{-4}$ | 3.09 | $3.68 \times 10^{-6}$ | |
| *NQM1* | Transaldolase of unknown function | 2.30 | $4.70 \times 10^{-3}$ | 3.47 | $4.60 \times 10^{-4}$ | Yes |
| *SOL4* | 6-phosphoglucono lactonase | 2.77 | $7.18 \times 10^{-5}$ | 2.46 | $1.10 \times 10^{-4}$ | |
| *TKL2* | Transketolase | 6.44 | $7.35 \times 10^{-7}$ | 2.62 | $1.19 \times 10^{-3}$ | |
| Stress protection and reserve carbohydrates |  |  |  |  |  |  |
| *GDB1* | Glycogen debranching enzyme (degradation) | 2.57 | $7.52 \times 10^{-3}$ | 3.66 | $3.40 \times 10^{-3}$ | |
| *GIP2* | Putative regulatory subunit of protein phosphatase Glc7 | 4.80 | $6.76 \times 10^{-8}$ | nd | nd | |
| *GLC3* | Glycogen branching enzyme (accumulation) | 2.89 | $5.33 \times 10^{-6}$ | nrc | nrc | |
| *GPH1* | Glycogen phosphorylase (mobilization) | 3.51 | $7.79 \times 10^{-5}$ | 3.13 | $6.20 \times 10^{-4}$ | Yes |
| *GSY1* | Glycogen synthase isoenzyme 1 | 3.07 | $6.68 \times 10^{-5}$ | nrc | nrc | |
| *GSY2* | Glycogen synthase isoenzyme 2 | 2.13 | $1.62 \times 10^{-3}$ | 2.70 | $1.30 \times 10^{-4}$ | Yes |
| *PGM2* | Phosphoglucomutase | 2.55 | $1.35 \times 10^{-3}$ | nrc | nrc | Yes |
| *TFS1* | Inhibitor of Ras GAP (Ira2p) and carboxy-peptidase Y (Prc1p) | 2.64 | $5.05 \times 10^{-5}$ | 3.00 | $1.06 \times 10^{-3}$ | Yes |
| *TPS2* | Phosphatase subunit of T-6-P synthase/phosphatase | 2.42 | $3.00 \times 10^{-4}$ | 2.50 | $3.90 \times 10^{-4}$ | Yes |
| *TSL1* | Large subunit of the T-6-P synthase/phosphatase | 2.48 | $1.36 \times 10^{-3}$ | 2.65 | $3.20 \times 10^{-4}$ | Yes |
| Cell surface and cell wall architecture |  |  |  |  |  |  |
| *CCW22* | Cell wall protein | 2.52 | $1.35 \times 10^{-30}$ | nrc | nrc | |
| *ECM4* | S-glutathionyl-(chloro) hydroquinone reductase | 2.03 | $8.96 \times 10^{-5}$ | 2.49 | $9.89 \times 10^{-3}$ | Yes |
| *LSP1* | Eisosome core component | nrc | nrc | 2.30 | $1.60 \times 10^{-4}$ | Yes |
| *PIR3* | Cell wall protein | 0.38 | $8.02 \times 10^{-6}$ | nd | nd | |
| *SCW10* | Cell wall protein | 0.45 | $2.61 \times 10^{-3}$ | nrc | nrc | |
| *YLR042C* | Cell wall protein | 0.42 | $1.88 \times 10^{-3}$ | nd | nd | |
| *YPS3* | Aspartic protease | 0.17 | $2.88 \times 10^{-20}$ | nd | nd | |
| *ZPS1* | Putative GPI-anchored protein | 3.87 | $7.81 \times 10^{-25}$ | nrc | nrc | |
| Mitochondrial functions |  |  |  |  |  |  |
| *ALD4* | Aldehyde dehydrogenase | nrc | nrc | 2.10 | $1.42 \times 10^{-3}$ | Yes |
| *CYB5* | Cytochrome b5 | 0.46 | $7.13 \times 10^{-4}$ | nrc | nrc | |
| *CYC1* | Cytochrome c, isoform 1 | 0.23 | $3.78 \times 10^{-13}$ | nrc | nrc | |
| *GUT2* | Glycerol-3-phosphate dehydrogenase | ns | ns | 2.21 | $1.36 \times 10^{-5}$ | |
| *NCA3* | Protein involved in mitochondrion organization | 0.34 | $5.10 \times 10^{-4}$ | nd | nd | |
| *NDE2* | External NADH dehydrogenase | 2.23 | $1.89 \times 10^{-2}$ | nd | nd | |
| *OM14* | Mitochondrial outer membrane receptor for cytosolic ribosomes | ns | ns | 2.08 | $1.94 \times 10^{-3}$ | |
| *OM45* | Mitochondrial outer membrane protein of unknown function | 2.96 | $8.08 \times 10^{-5}$ | 3.44 | $5.60 \times 10^{-4}$ | |
| *PIC2* | Copper/phosphate carrier | 2.00 | $2.44 \times 10^{-6}$ | nd | nd | |
| *SFC1* | Succinate-fumarate transporter | 2.11 | $2.66 \times 10^{-2}$ | nd | nd | |
| *YMC2* | Putative mitochondrial inner membrane transporter | 0.46 | $2.37 \times 10^{-3}$ | nrc | nrc | |

nrc = no relevant changes; nd = not detected; T-6-P = trehalose-6-phosphate
<sup>a</sup> Due to high sequence similarity, these yeast hexose transporters cannot be differentiated from the peptides detected in mass spectrometry. Data could thus reflect the concentration changes in either one of these transporters or both.
<sup>b</sup> Where stress responsive elements (STREs) have either been shown to function in gene expression, or at least were detected in bioinformatic surveys (Aranda and del Olmo MI, 2003; Chautard et al., 2004; Francois et al., 1992; Moskvina et al., 1998; Parrou et al., 1997; Verwaal et al., 2004; Winderickx et al., 1996), their presence is indicated as "yes".

Under oxidative stress, i.e. exposure to 0.8 mM H_2_O_2_ for six hours, candidate genes involved in the general or environmental stress response (ESR) were upregulated both in the wild-type and the *rho5* deletion strains (Fig. 2B). Differential regulation between the two genetic backgrounds was observed for 120 genes (upregulation) and for 276 genes (downregulation; supplementary Table S2).

In a second approach, differential protein concentrations for the same strains under standard growth conditions were assessed by mass spectrometry. A total of 48 proteins increased significantly in their amount when the deletion mutant was compared to the wild type, whereas only 13 proteins showed a decreased concentration (Fig. 2C; with cut-offs applied at p-values less than 0.05 and at least a twofold change in protein abundance; see supplementary Table S3 for a complete list of affected proteins). Again, proteins most strongly affected by the *rho5* deletion included those involved in carbohydrate metabolism (Table 1), several overlapping with and confirming the RNAseq data (Fig. 3).

**Figure 3.**
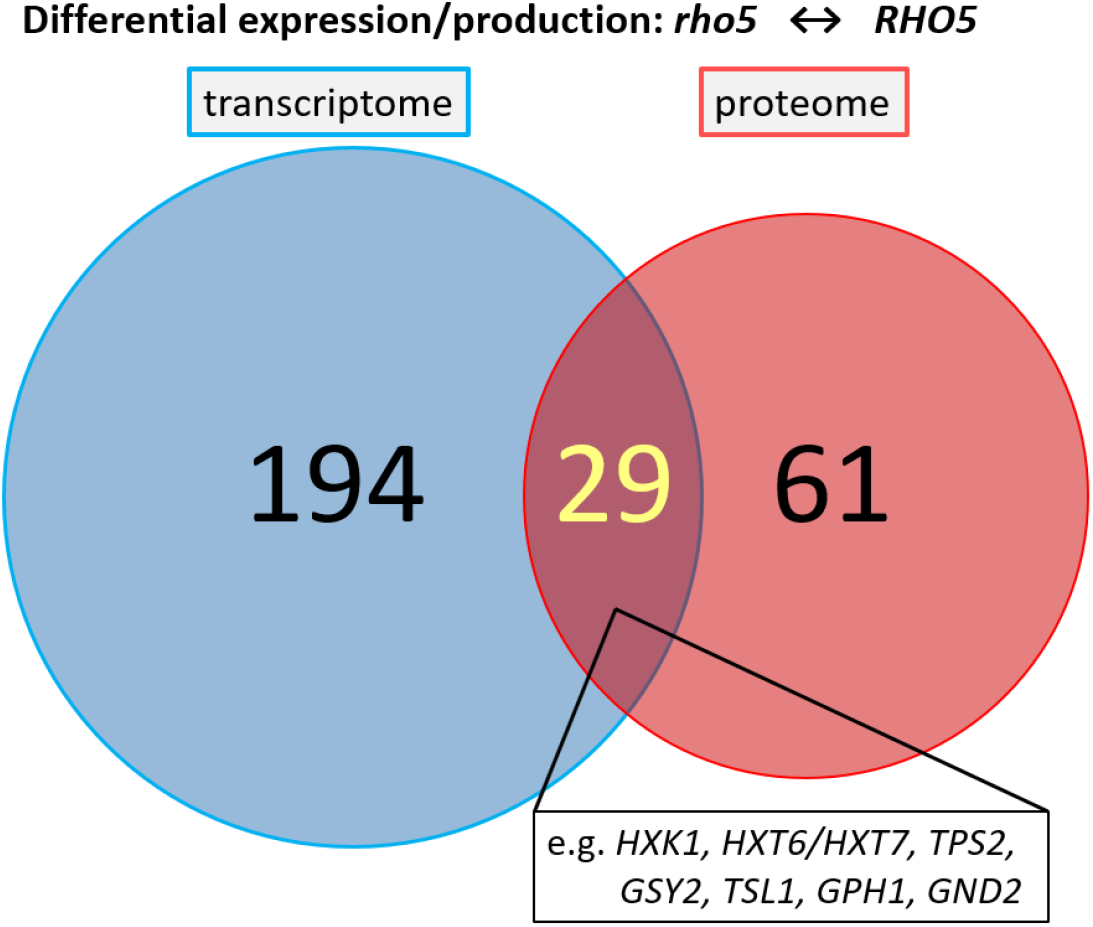
Venn-Diagram of overlapping data obtained from RNA-sequencing and mass spectrometry from comparison of a *rho5* deletion (FSO62-7A) to a wild-type strain (HD56-5A) after growth on SCD. Of the 29 overlapping results, 27 genes/ proteins were up- and two were down-regulated. Some examples of components related to carbohydrate metabolism are indicated. Cut-off critera for significant differences were the same as described in the legend of Figure 2.

### 2.2. Epistasis analyses reveal genetic interactions of RHO5 with glucose signaling through the SNF1 complex and hexokinase

The global expression data suggested a relationship between Rho5 and carbohydrate metabolism. We therefore proceeded by assessing the phenotypes of either a *rho5* deletion or the hyper-active *RHO5^G12V^* allele in combination with different mutants in the major glucose signaling pathways. For this purpose, classical genetic crosses were performed, the resulting diploids were subjected to tetrad analyses, and growth was first monitored by determination of colony sizes of the different mutant and wild-type segregants on rich medium plates with 2% glucose as a carbon source.

As quantified in Fig. 4A, segregants with a *rho5* deletion form only slightly smaller colonies than those with the wild-type allele. However, the growth area is reduced by approximately threefold in segregants lacking either the kinase subunit of the SNF1 complex (*snf1*) or the Reg1 subunit of its phosphatase, which is required for its inactivation (note that in lack of a hyper-active Snf1 kinase derivative *reg1* deletions are commonly employed to constitutively activate the SNF1 complex; Ludin et al., 1998). Interestingly, an additional *rho5* deletion aggravates the growth defect of strains lacking Reg1, while it does not alter growth of the *snf1* deletions (Fig. 4A). The fact that the slow growth of *rho5 reg1* strains is restored back to that of a single *reg1* deletion by an additional lack of Snf1, i.e. in *rho5 reg1 snf1* triple deletions, suggests that Rho5 negatively affects the activity of the SNF1 complex. These growth impairments derived from colony sizes were also confirmed by recording growth curves in liquid synthetic medium, suggesting that they are indeed owed to differential growth rates, rather than a delay in spore germination (Fig. S1A).

**Figure 4.**
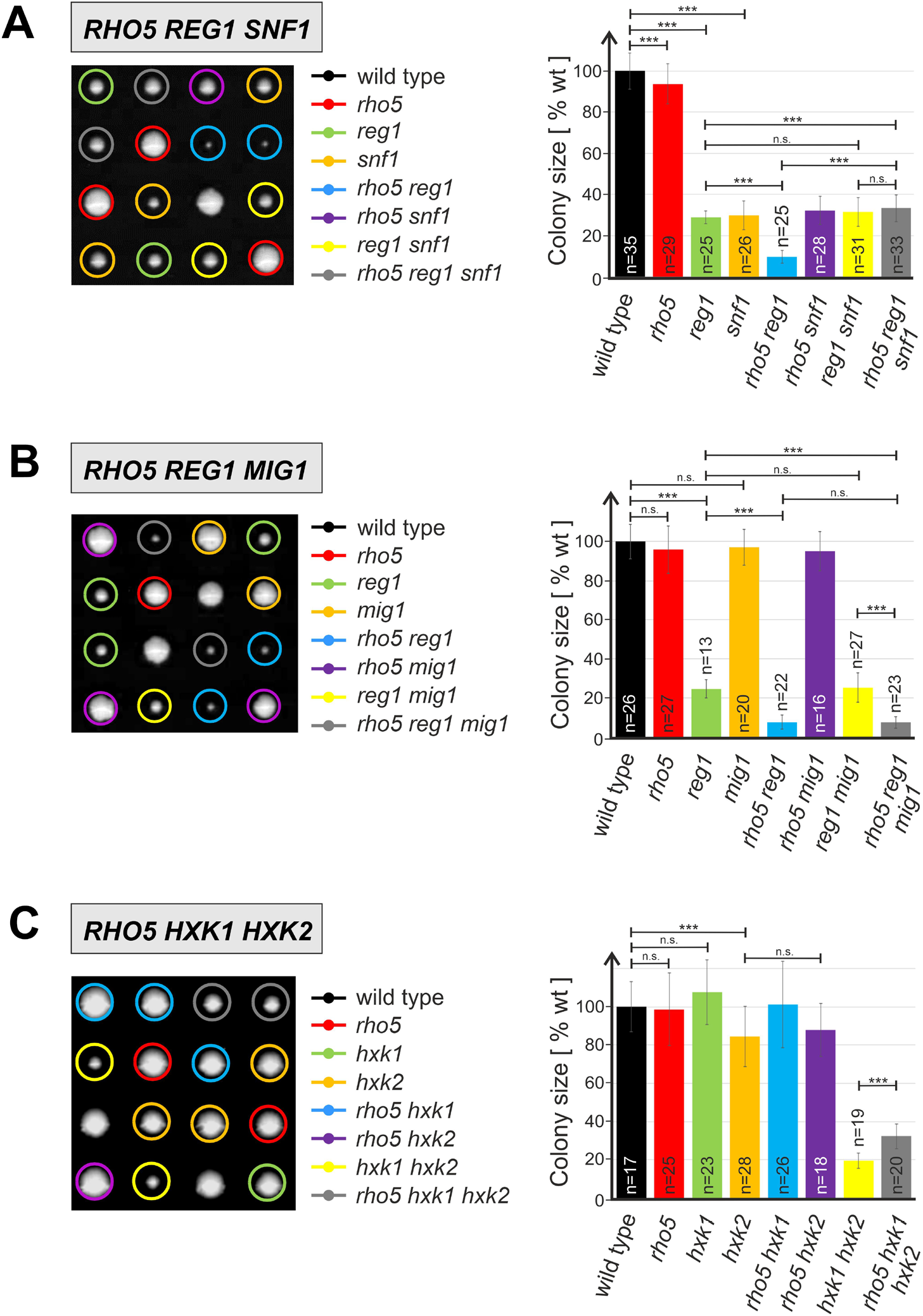
Epistasis analyses based on growth of segregants from tetrad analyses on rich medium (YEPD). Plates were incubated for three to five days at 30 °C, depending on the crosses to be analyzed. Only four exemplary tetrads are shown for each cross, with colored circles designating different combinations of gene deletions as indicated. Colony sizes for each combination (determined from pixel area and given as percentage from wild type, which was set at 100%) were determined from at least 40 tetrads from each cross and quantified in the columns of the diagram at the right (n = total number of segregants obtained for each genotype; error bars are indicated for each data set, three asterisks indicate highly significant differences with p-values below 0.001; n.s. = not significant).

Diploids analyzed were obtained from the following crosses: A) A strain carrying a *rho5 snf1* double deletion (FSO67-2C) with a *reg1 snf1* double deletion (FSO66-3C). B) A strain carrying a *rho5* deletion (FSO71-2A) with a *reg1 mig1* double deletion strain (FSO79-8C). C) A strain carrying a *rho5* deletion (FSO43-1D) with one carrying a *hxk1 hxk2* double deletion (HOD257-2B).

As the transcription factor Mig1 is a major downstream target of SNF1 signaling in carbohydrate metabolism, epistatic relationships were also investigated with a *mig1* deletion. Surprisingly, the growth retardation observed in a *rho5 reg1* double deletion, amounting to less than 10% of wild-type segregants, was not alleviated in a triple *rho5 reg1 mig1* deletion (Figs. 4B and supplementary Fig. S1B), demonstrating that the observed Rho5- and Snf1-dependent growth effects are not mediated by Mig1. In contrast, the growth defect caused by a lack of Snf1 requires a functional Mig1, as it is relieved in strains with a *snf1 mig1* double deletion (Fig. S2).

In parallel to the SNF1 complex, hexokinase PII was among the first components found to participate in yeast glucose repression (Entian, 1980; Entian and Zimmermann, 1980). Besides its association with Mig1 in the nucleus at high external glucose concentrations, the inhibitory action of Hxk2 on the SNF1 complex is probably associated with its catalytic activity, but still somewhat enigmatic (reviewed in (Gancedo, 2008). *HXK2* encodes one of three yeast isozymes capable of glucose phosphorylation, together with a glucokinase encoded by *GLK1* and another hexokinase encoded by *HXK1*. Expression of the latter two is subject to glucose repression and only *hxk1 hxk2 glk1* triple deletions cannot grow on glucose as a sole carbon source (Rodriguez et al., 2001; Walsh et al., 1991). As expected, we found only a moderate decrease in colony sizes after tetrad analyses for *hxk2* deletions compared to wild-type segregants, whereas those of *hxk1 hxk2* double deletions were reduced by approximately 80% (Fig. 4C). Interestingly, this phenotype could be partially alleviated by an additional *rho5* deletion, which restored growth of the triple mutants to approximately 30% of that of the wildtype colonies. Again, these findings were substantiated by recording growth curves of segregants with the different mutant combinations (Fig. S1C). We attribute the slight positive effect of the *rho5* mutants to the increase in respiratory capacity, as reported below.

### 2.3. *RHO5* genetically interacts with cAMP/PKA signaling

Next, we addressed the relationship between Rho5 and the cAMP/PKA signaling pathway. Since growth of mutants lacking components of that pathway is generally not impaired under standard growth conditions, we employed their sensitivity towards hydrogen peroxide as a readout for epistasis analyses with *rho5* deletions. Although PKA also phosphorylates several cytosolic enzymes involved in carbohydrate metabolism, its major effect on nuclear gene expression at high glucose concentrations is the inactivation of the redundant transcription activators Msn2/Msn4, by triggering their nuclear export (Fig. 1; reviewed in Busti et al., 2010; Heinisch and Rodicio, 2017). Since these transcription factors also mediate the yeast’s general stress response, it is not surprising that *msn2 msn4* double deletions display an increased sensitivity towards oxidative stress exerted by hydrogen peroxide (Fig. 5A). This phenotype cannot be rescued by an additional *rho5* deletion, which on its own shows hyper-resistance, indicating that Rho5 acts upstream of Msn2/Msn4 in the signaling cascade.

**Figure 5.**
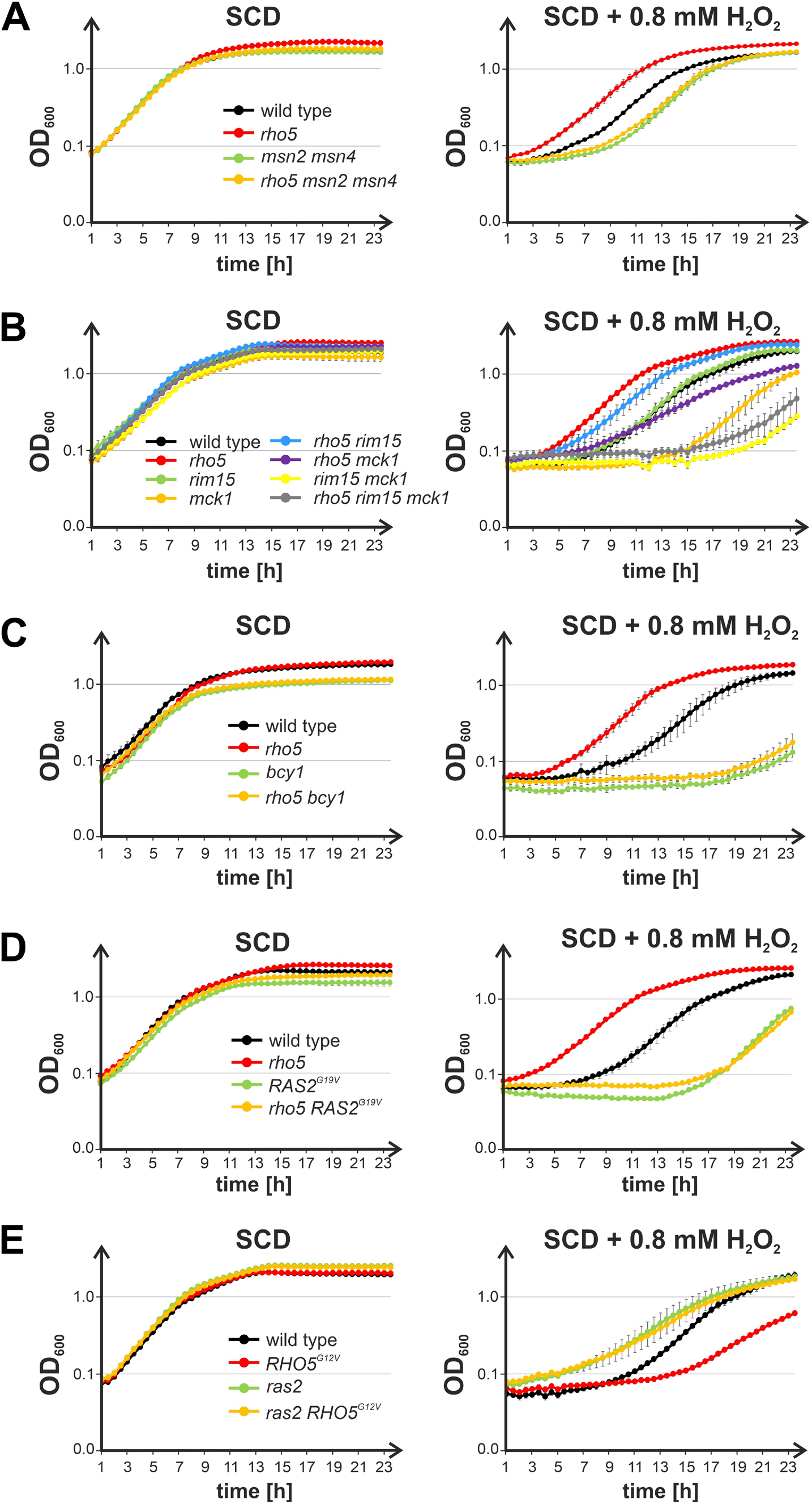
Epistasis analyses based on the sensitivity towards hydrogen peroxide of strains carrying mutations in genes encoding components of the cAMP/PKA signaling pathway in combination with *RHO5* variants.

Growth curves were recorded on synthetic medium with 2% glucose (SCD; supplemented with 4 mg/L histidine for strains with a histidine auxotrophy), in the absence and presence of hydrogen peroxide as indicated. Error bars give the standard deviations at each time point obtained from at least two biological and two technical replicates from parallel measurements for each curve (i.e. two independent isogenic segregants were measured, with two independent inoculates, each). Genotypes of the otherwise isogenic strains are listed in Table 2. Strains employed were: A) Wild type (FSO36-5B and FSO36-11A), *rho5* (FSO36-11D and FSO36-5D), *msn2/4* (FSO36-11B and FSO36-5A), *rho5 msn2/4* (FSO36-5C and FSO36-11C). B) Wild type (FSO86-7A and FSO86-2A), *rho5* (FSO105-7D and FSO105-1C), *rim15* (FSO86-6B and FSO86-1C), *mck1* (HOD343-2A and HOD343-4A), *rho5 rim15* (FSO86-7C and FSO86-2C), *rho5 mck1* (HOD343-2C and HOD343-4C), *rim15 mck1* (FSO84-4A and FSO84-10B), *rho5 rim15 mck1* (FSO84-10D and FSO84-3D). C) Wild type (FSO16-3B and FSO86-7A), *rho5* (FSO105-7D and FSO105-1C), *bcy1* (FSO105-4A and FSO105-1B), *rho5 bcy1* (FSO105-1D and FSO105-7C). D) Wild type (FSO35-4A and FSO36-5B), *rho5* deletion (FSO36-11D and FSO105-7D), *RAS2^G19V^* mutant strain (HOD610-1D/RAS2^G19V^URA and HOD610-2C/RAS2^G19V^URA) and a *rho5 RAS2G^19V^* mutant strain (HODrk21 and HODrk22). E) Wild type (FSO35-2A and FSO35-4A), *RHO5^G12V^* (FSO88-3A and FSO88-2B), *ras2* (FSO98-6B and FSO98-3A), *RHO5^G12V^ ras2* (FSO78-14B and FSO78-6C).

**Table 2.**
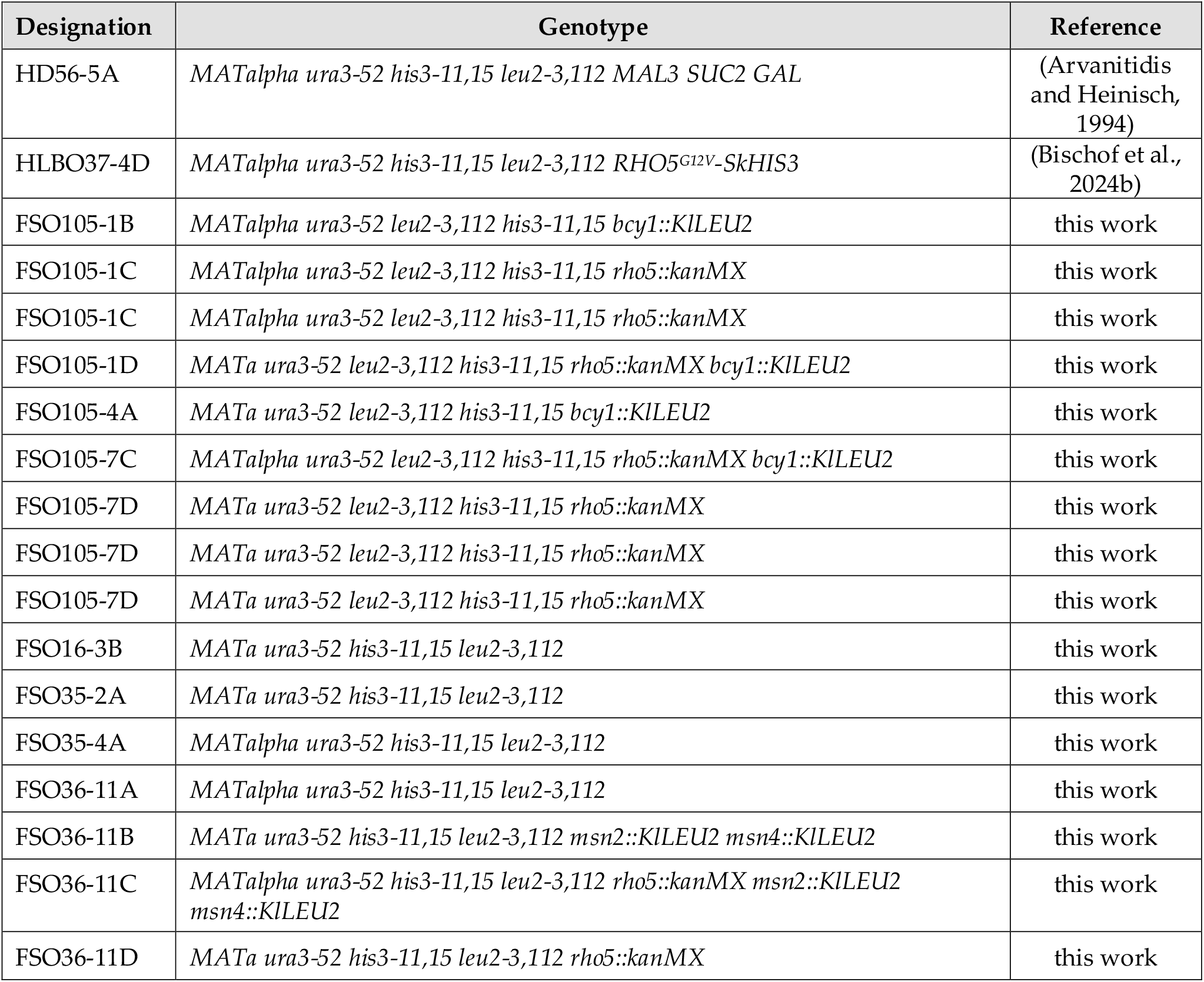
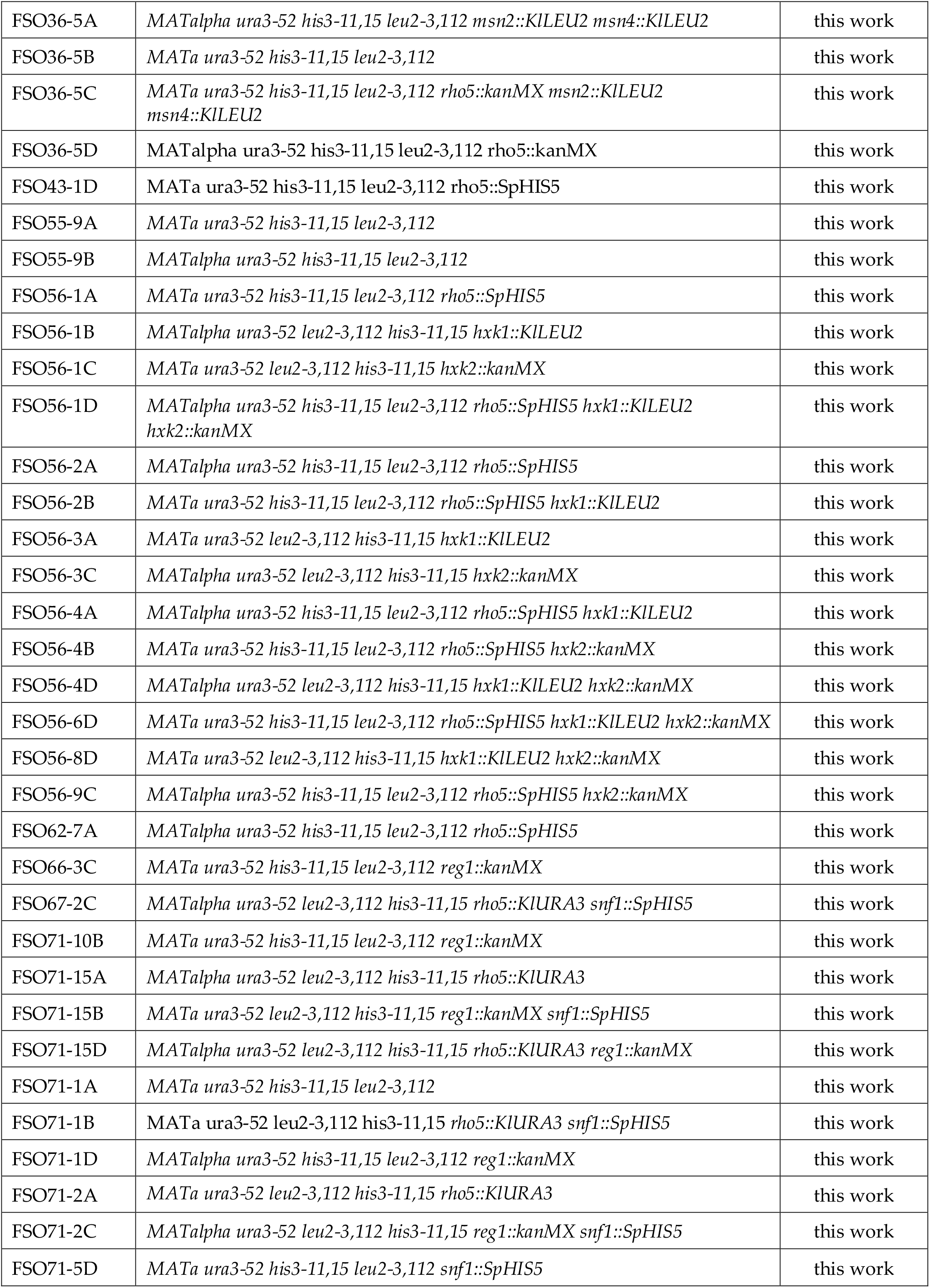
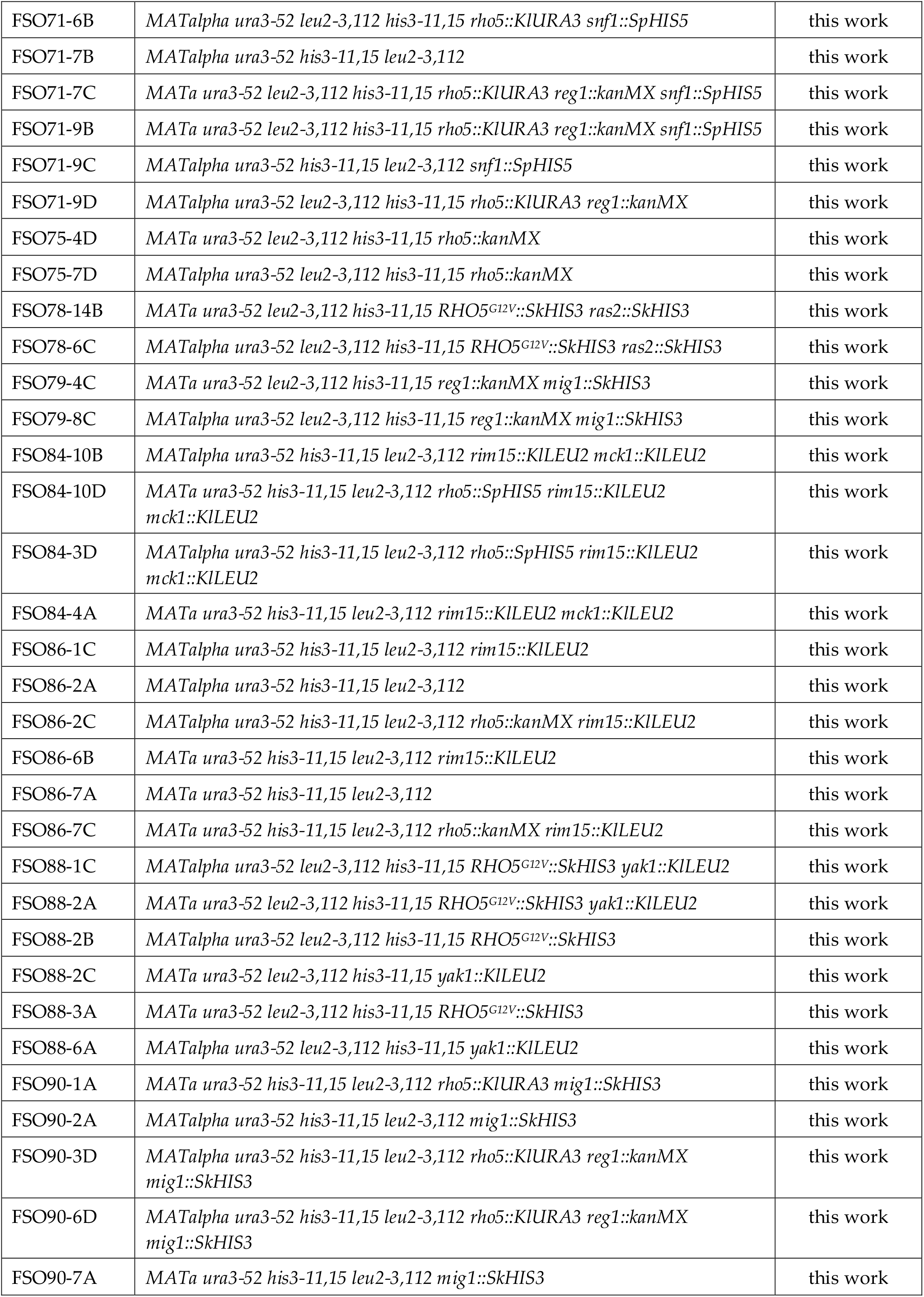
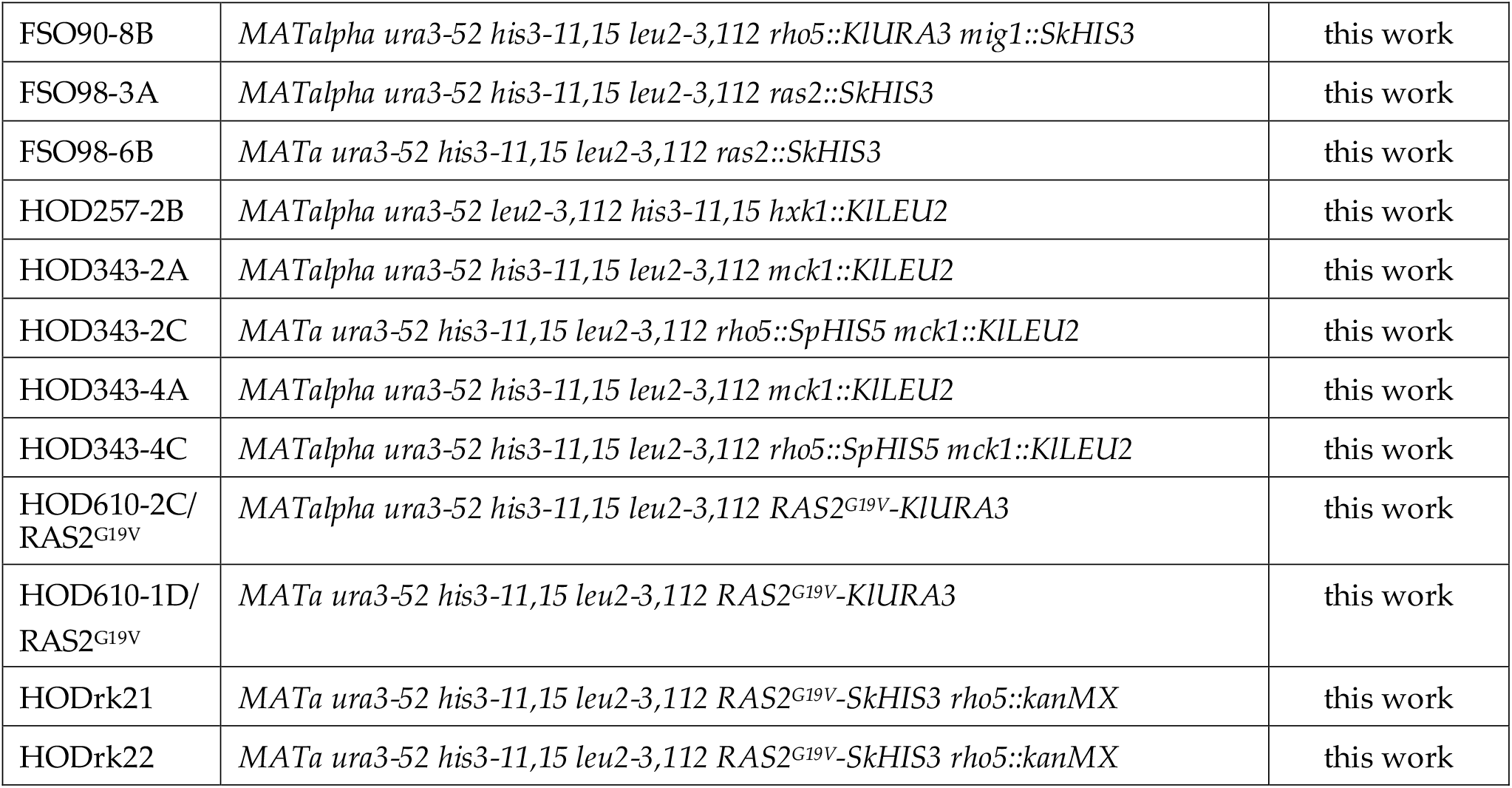
Yeast strains employed in this work

To further identify at which stage Rho5 interferes with the signaling cascade, we consecutively deleted genes encoding other upstream components. Besides being a direct target for inhibition by PKA-mediated phosphorylation, Msn2/Msn4 can be activated by the protein kinases Rim15, Mck1, and Yak1, which are inhibited by PKA-mediated phosphorylation (Cao et al., 2016). Growth curves of the respective mutants revealed that strains lacking Yak1 grew similar to wild-type in the presence and absence of hydrogen peroxide, indicating that this kinase does not mediate Rho5-dependent growth effects (supplementary Fig. S3). Yet, a *rim15* deletion caused a moderate increase in sensitivity towards hydrogen peroxide as compared to wild-type cells, whereas *mck1* deletions were clearly hyper-sensitive, with an additive effect in the *rim15 mck1* double deletions (Fig. 5B). An additional lack of Rho5, which by itself causes a distinct hyper-resistance towards the stressor, did not lead to a pronounced improvement of growth of the triple *rim15 mck1 rho5* mutants under oxidative stress. This suggests that Rho5 also acts upstream of, but clearly through, the two kinases.

In the presence of glucose and under standard growth conditions the protein kinase A subunits (Tpk1, Tpk2, Tpk3), placed further upstream in the cAMP/PKA signaling cascade, dissociate from the regulatory subunits encoded by *BCY1* and the heterotetramer, so that the kinase subunits can phosphorylate their target proteins. We thus constructed *bcy1* deletions to obtain strains with a constitutively active PKA. These displayed a pronounced hyper-sensitivity towards hydrogen peroxide, which could not be rescued by an additional *rho5* deletion (Fig. 5C).

A major component activating the yeast adenylate cyclase and thereby PKA is another small GTPase, Ras2. To confirm the results obtained with the *bcy1* deletion, we therefore expressed a hyperactive *RAS2^G19V^* allele, which caused an increased sensitivity towards hydrogen peroxide compared to the wild type, again not influenced by the presence or absence of Rho5 (Fig. 5D). *Vice versa*, a *ras2* deletion clearly led to an increase in oxidative stress resistance. The latter was not suppressed by expression of a hyper-active *RHO5^G12V^* allele, which leads to a hyper-sensitive phenotype by itself (Fig. 5E). Taken together, these data demonstrate that Rho5 acts upstream of the cAMP-dependent PKA and Ras2 in the yeast’s nutrient and general stress response.

### 2.4. Rho5 affects respiratory functions

As *S. cerevisiae* is a Crabtree-positive yeast, respiration is significantly inhibited in the presence of 2% glucose in the medium (Lagunas et al., 1982). Growth may thus be partially limited by a reduced energy supply and the increase in colony sizes after tetrad analysis in the *rho5* deletion background observed above, e.g. in the triple deletion with *hxk1* and *hxk2*, could be a result of increased respiratory capacity. We therefore determined the effect of either the lack or the hyper-activation of Rho5 on respiration when growing the strains on 2% glucose medium. As evident from Fig. 6, lack of Rho5 leads to a duplication in the rate of respiration as compared to the wild-type, whereas the hyper-active Rho5^G12V^ variant caused only a minor, less significant increase. We conclude that Rho5 has some inhibitory effect on the rate of respiration, although how this is exactly mediated remains to be determined (see discussion).

**Figure 6.**
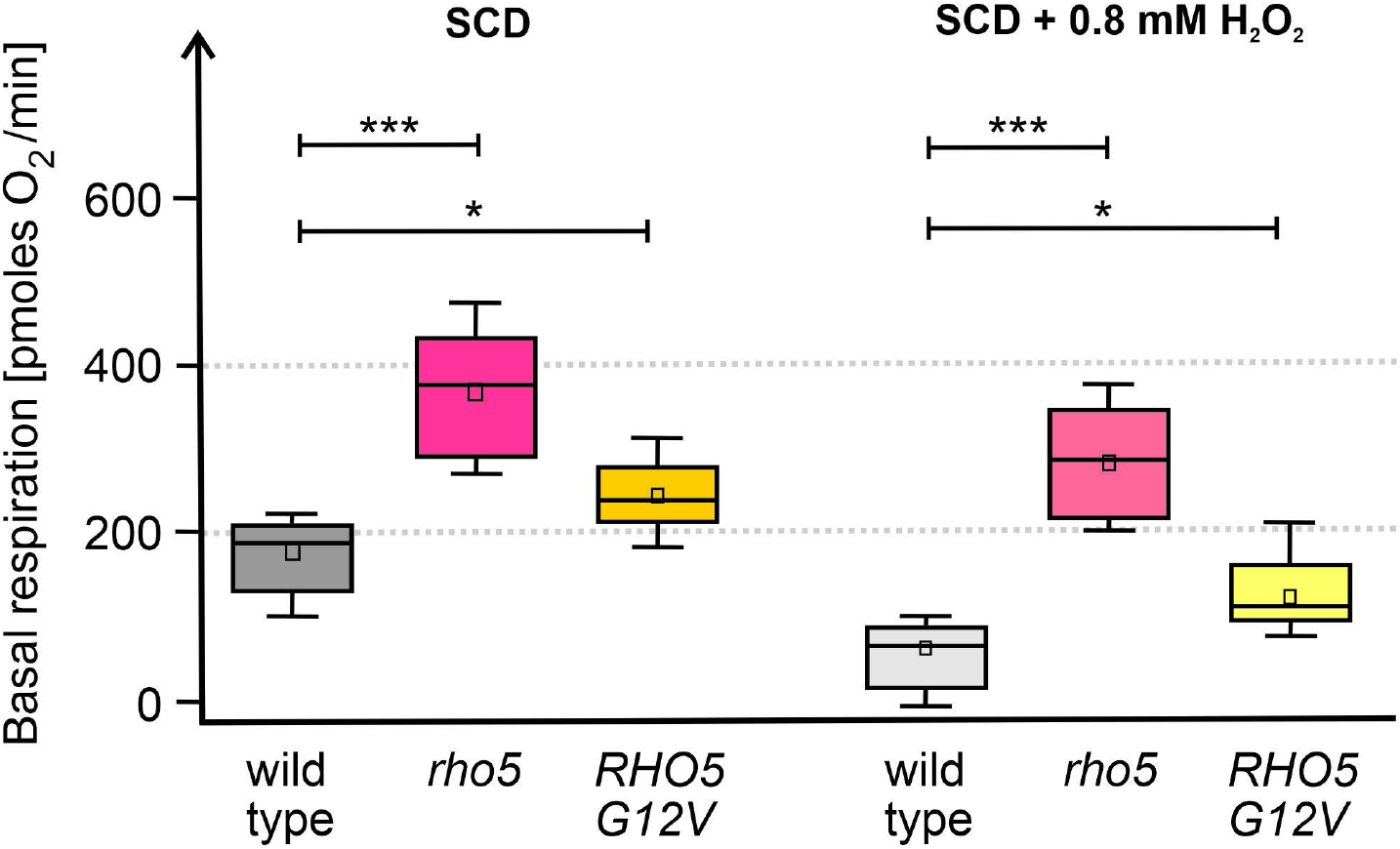
Respiration measurement of a wild-type strain (HD56-5A), a *rho5* deletion strain (FSO62-7A) and a *RHO5^G12V^* mutant strain (HLBO37-4D). Significance is indicated by one asterisk, while three asterisks indicate a very high significance. Note that differences between each strain with and without hydrogen peroxide are also highly significant but not shown here for the sake of clarity.

## 3. Discussion

Following previous hints for a participation of the small yeast GTPase Rho5 in the response to nutrient availability (Schmitz et al., 2018) a more detailed analysis of its crosstalk with the known glucose signaling pathways was initiated in this work. To substantiate the physiological relevance of the Rho5 molecular switch, we started by global analyses of both the transcriptome and the proteome in a *rho5* deletion *versus* a wild-type strain, which will first be discussed.

### 3.1. Gobal expression analyses

Given that Rho5 was first described as a negative regulator of cell wall integrity (CWI) signaling (Schmitz et al., 2002), it was not surprising to find the upregulation of genes and/or proteins that influence the architecture of the cell surface in a *rho5* deletion. This extends to eisosomal proteins, as their turnover was shown to be mediated by Slt2, the downstream mitogen-activated protein kinase (MAPK) of the CWI pathway (Mascaraque et al., 2013). Likewise, an increase in the concentration of some mitochondrial proteins and/or the expression of their encoding genes was detected. This can be explained by the participation of wild-type Rho5 in stress-dependent mitophagy, resulting in a reduced rate of mitophagy in the deletion mutant, which should affect retrograde signaling for nuclear gene expression (Bui and Labedzka-Dmoch, 2024; Jazwinski, 2014). On the other hand, the downregulation of genes encoding other mitochondrial proteins such as Cyb5 and Cyc1 in the *rho5* deletion indicates that the observed increase in respiratory activity may be mediated by post-translational effects.

Of major interest in the context of this work was the identification of several genes and proteins involved in carbohydrate metabolism that were upregulated in the absence of Rho5. Thus, several hexose transporters, encoded by *HXT* genes, increased in gene expression, protein concentration, or both. These comprised the high-affinity transporters Hxt6/Hxt7, whose almost identical sequences prohibit their differential analysis by mass spectrometry (Reifenberger et al., 1995), and Hxt5 with a moderately high affinity for glucose. All three genes are expressed at low glucose concentrations, as confirmed by transcriptome analyses in wine fermentations (Boles and Hollenberg, 1997; Perez et al., 2005). In contrast, the expression of the *HXT11* gene, encoding another member of the transporter family, was found to be significantly downregulated in a *rho5* deletion. This could be due to the role of this transporter in pleiotropic drug resistance, and to a different set of transcription factors, Pdr1 and Pdr3, which control its gene expression (Nourani et al., 1997). The gene encoding the hexokinase isoform 1, *HXK1*, subject to repression at high glucose concentrations as discussed for the *HXT* genes (Rodriguez et al., 2001), was also upregulated both at the transcriptional and the protein level in a *rho5* deletion. This suggested a general role of Rho5 in glucose repression which was supported by our data on epistatic relationships and respiratory capacity, as discussed below.

Regarding the pentose phosphate pathway (PPP), it is believed to contribute only to a minor extent to glucose metabolism in *S. cerevisiae* as compared to other yeasts (reviewed in Bertels et al., 2021). Nevertheless, its oxidative part, especially the reaction of glucose-6-phosphate dehydrogenase (Zwf1), produces reduction power (NADPH) required to cope with oxidative stress (Campbell et al., 2016; Heinisch et al., 2020; Thomas et al., 1991). Rho5 also contributes to a proper response to such stress (Schmitz et al., 2015; Singh et al., 2008; Singh et al., 2019). Interestingly, a lack of Rho5 leads to an increased expression of the genes encoding some of the apparently less relevant isozymes acting in the pathway, like *SOL4*, *GND2*, and *TKL2* (Juhnke et al., 1996; Schaaff-Gerstenschlager et al., 1993; Stanford et al., 2004). In this context, *NQM1*, which encodes a putative transaldolase isozyme with a low catalytic activity (Huang et al., 2008), is repressed by glucose, and was proposed to regulate transcription of genes involved in stress resistance in stationary phase cells (Michel et al., 2015). One could thus assume that while the major isozymes of the PPP catalyze the housekeeping reactions, alternative isozymes are upregulated to cope with stressful conditions such as glucose depletion and the presence of hydrogen peroxide, a response involving the Rho5 molecular switch.

Finally, we found that genes and enzymes related to the biosynthesis of the stress-protectant disaccharide trehalose and the turnover of the reserve polysaccharide glycogen, both in its synthesis and its degradation, are upregulated in a *rho5* deletion as compared to a wild-type strain (see Francois and Parrou, 2001, for a comprehensive review on yeast reserve carbohydrate metabolism). Both compounds accumulate while yeast cells are approaching stationary phase from a glucose-rich medium (Lillie and Pringle, 1980), and were found to be significantly reduced in cells lacking Rho5 (Cao et al., 2016). The upregulation of these genes in a *rho5* deletion could be expected, as they are controlled by Msn2 and Msn4 through the STRE binding sites in their promoters. That the accumulation of glycogen and trehalose depends on the specific stress applied and does not always coincide with the level of gene expression for the enzymes involved in their synthesis has been noted, before (Francois and Parrou, 2001). Nevertheless, our findings underline the previously proposed important role of Rho5 in regulating the cells response to both nutrient limitation and oxidative stress (Schmitz et al., 2018) and are reflected in the transcriptome and proteome data.

### 3.2. Interactions of Rho5 with Hxk2- and SNF1-dependent glucose signaling

The apparent upregulation of many hexose transporters and genes subject to glucose repression observed in the transcriptome and proteome analyses prompted a deeper analysis of the genetic interactions of *RHO5* with genes encoding components in the known glucose signaling pathways depicted in Fig. 1. As expected from previous reports, *hxk2* null mutants displayed only a mild growth defect, attributed to the fact that yeast has two further kinases capable of glucose phosphorylation, Hxk1 and Glk1 (Walsh et al., 1991). Although the latter are usually not expressed in glucose media due to catabolite repression, this regulation is alleviated by a lack of Hxk2, thus restoring the ability to channel glucose into glycolysis (Entian, 1980; Rodriguez et al., 2001). Glucose signaling is exerted by Hxk2 as a moonlighting enzyme that can enter the nucleus and act as a coactivator of the Mig1 repressor (reviewed in Gancedo, 2008). Since the gene encoding glucokinase, *GLK1*, also carries a STRE in its promoter, the improved growth of a *hxk1 hxk2* double deletion by an additional lack of Rho5 could be attributed to its increased expression mediated by Msn2 and Msn4. The GTPase may also have a secondary effect: The number of mitochondria and cellular respiration are controlled by glucose repression in *S. cerevisiae* (Böker-Schmitt et al., 1982; Lagunas et al., 1982). Apparently, wildtype Rho5 contributes to this repression, as it is partially alleviated in the *rho5* deletion. The limited amount of glucose phosphorylated by the remaining glucokinase in the triple deletion strains could then be more efficiently used for ATP generation by an increased rate of respiration in the *rho5* null mutant compared to the wild-type, as found herein. We believe that the influence of Rho5 on respiration may be rather indirect, as the deletion mutant shows a significant increase, but the hyper-active Rho5^G12V^ variant has no diminishing effect, as would be expected from a direct regulation. Thus, the reduced mitophagy observed in *rho5* deletions (Schmitz et al., 2015; Singh et al., 2008) may result in the presence of more mitochondria and respiratory chain components, leading to an enhanced ATP supply.

With regard to signaling through the SNF1 complex we found that a lack of the kinase activity causes a growth defect independent of Rho5. On the other hand, hyper-activation of the complex, achieved by deletion of the gene encoding the inhibitory Reg1 phosphatase, also severely impaired growth, but was aggravated by a lack of Rho5. Further epistasis analyses then showed that the slow growth a *reg1* deletion depended on the upregulation of the SNF1 pathway, which commonly is mediated by the transcriptional repressor Mig1 (Fig. 1; Hedbacker and Carlson, 2008). However, the genetic interaction between *RHO5* and *REG1* required only SNF1 activity, but not Mig1. This indicates that other targets of SNF1 are mediating the Rho5-dependent response, with the redundant transcription factors Msn2/Msn4 being primary candidates, as discussed below in the interaction with PKA signaling. It should be noted that an indirect effect of Rho5 on growth by its signaling to cell wall synthesis through SNF1 can be excluded, as this would also be mediated by Mig1 (Backhaus et al., 2013; Rippert et al., 2017).

### 3.3. Interactions of Rho5 with cAMP/PKA signaling

To facilitate the epistasis analyses between *rho5* and mutants defective in cAMP/PKA signaling we employed their effect on oxidative stress response: While strains lacking Rho5 are more resistant towards hydrogen peroxide than wild type, blocks in the cAMP/PKA pathway cause an increased sensitivity (Hasan et al., 2002; Schmitz et al., 2015; Singh et al., 2008). The latter would be expected to be rescued in double mutants with *rho5* if the GTPase acted in a downstream step of the signaling pathway or in a parallel route. On the other hand, cells would be expected to remain hyper-sensitive if Rho5 was acting upstream of the block in the same signaling pathway. Moreover, opposite results would be expected when hyper-active mutant alleles were employed in the epistasis analyses. For example, the hyper-sensitivity of a Rho5^G12V^ variant towards hydrogen peroxide would be rescued by blocking a step downstream in the same, but not in a different, signaling pathway. Under these premisses, our results clearly show that *RHO5* is epistatic to *RAS2* and all other genes encoding downstream components of the cAMP/PKA pathway depicted in Fig.1.

Interestingly, the signal to PKA elicited by Rho5 under oxidative stress seems to be transmitted to the transcription factors Msn2/Msn4 predominantly by the branch inactivating the Rim15/Mck1 kinases, as the hyper-sensitivity of the double deletion mutants cannot be rescued by a *rho5* deletion. Note that the degree of resistance mediated by the two kinases is additive, with our data suggesting that Mck1 plays a more prominent role in this case. These findings are in line with previous reports indicating that the two kinases mediate Rho5-dependent survival under glucose starvation (Cao et al., 2016). Similar to our results, Rim15, but not a third protein kinase Yak1, was also found to differentially control gene expression of *TPK1*, encoding one of the three PKA catalytic subunits, under heat shock and salt stress (Pautasso and Rossi, 2014). Moreover, we previously reported that a *rho5* deletion displays severe synthetic growth defects not only with *gpr1* and *gpa2* deletions affecting the other branch of glucose signaling to PKA (Fig. 1), but also with a *sch9* deletion (Schmitz et al., 2018). The latter lacks a protein kinase required to mediate nutrient signaling by TORC1, which inhibits Rim15 by Sch9-dependent phosphorylation, and thus integrates general nutrient with glucose signaling to fine-tune the activity of Msn2/Msn4 and thereby the expression of their target genes (Dawes and Perrone, 2020; Deprez et al., 2018). A feedback regulatory loop is then provided by Tps2, a trehalose-6-phosphate phosphatase, whose gene expression is under the control of Msn2/Msn4. *TPS2* expression was found to be upregulated in a *rho5* deletion in our RNAseq data and confirmed by the mass-spectrometry data. Tps2 mediates the dephosphorylation and activation of Rim15 by triggering its dissociation from Bmh1/Bmh2 thus facilitating its nuclear translocation (Kim et al., 2021).

### 3.4. To bind it all – a hypothesis for the role of Rho5 in glucose signaling

The data presented in this work and discussed above strongly indicate that the function of Rho5 in transmission of the glucose availability in the medium into a cellular response is largely mediated by PKA-dependent signaling to the transcription factors Msn2/Msn4. We pinpointed Rho5 to activate a step upstream of Ras2, with further signaling executed via the Rim15/Mck1 branch of protein kinases (Fig. 1). This holds true at least for the parallel response to oxidative stress, based on the epistasis analyses with deletion mutants within the cAMP/PKA pathway. As stress-related genes show an increased expression in cells lacking Rho5, the GTPase should activate the pathway in wild-type cells under standard growth conditions in glucose-rich media. How does this fit with the general expression analyses and the results on epistasis analyses for hexokinase and the SNF1 complex depicted in the left-hand part of Fig. 1?

The transcription factors Msn2/Msn4 are not only a target of phosphorylation for PKA, but in fact have first been identified as multicopy suppressors of a temperature-sensitive *snf1* mutant, as they are also inactivated by SNF1-dependent phosphorylation (Estruch and Carlson, 1993). Moreover, an inverse relationship was found for SNF1 and PKA activities, as one inhibits the other (reviewed in (Sanz et al., 2016). Both, a lack of kinase activity in the *snf1* deletion, as well as a constitutively high activity in the *reg1* deletion, reduce the ability to grow on glucose, as reported earlier in other strain backgrounds (Correa-Romero et al., 2023; Ruiz et al., 2011). We found this growth retardation to be aggravated in a *reg1 rho5* double deletion, which could be explained by a further increase in SNF1 kinase activity caused by the lack of the feedback inhibition between PKA and SNF1. Consequently, Msn2/Msn4-dependent gene expression would be strongly inhibited in a manner independent of Mig1, as observed. In contrast, the growth retardation observed in single *snf1* mutants and the *snf1 rho5* double deletion strains may be attributed to deregulation of Mig1-mediated repression.

Finally, the regulation through PKA and SNF1 also serves to embed Rho5 action in mitochondrial turnover and thereby in energy metabolism. We recently reported on the genetic interaction between *RHO5* and *POR1*, a gene encoding the major voltage-dependent anion channel (VDAC) in *S. cerevisiae* (Bischof et al., 2024b). In turn, the VDAC is required to regulate the activity and nuclear localization of the SNF1 complex (Shevade et al., 2018; Strogolova et al., 2012). Autophagy, and specifically mitophagy, are also interconnected at multiple levels with these signaling pathways. For instance, PKA inhibits autophagy by phosphorylation of Msn2/Msn4 (Yorimitsu et al., 2007). This function is coordinated with the response to other nutrient limitations mediated by the TORC1 complex and the protein kinase Sch9 and its activation of Rim15 (Yorimitsu et al., 2007). Interestingly, a *rho5* deletion is synthetically lethal with *sch9* null mutants, also indicating a concerted action of Rho5 through PKA and TORC1 signaling (Schmitz et al., 2018). And the activated SNF1 complex under glucose limitation also activates TORC1, ultimately promoting expression of Msn2/Msn4-dependent genes (Caligaris et al., 2023). These include *ATG8* and *ATG39*, which encode components of the autophagic machinery (Mizuno and Irie, 2021; Vlahakis et al., 2017). Another autophagy component, Atg21, was shown to interact directly with Rho5 (Singh et al., 2019). As Rho5 rapidly translocates to mitochondria under oxidative stress, it has been proposed to directly trigger mitophagy (Bischof et al., 2024b; Bischof et al., 2022). Whether or not this is related to its effect on Ras2 observed herein remains to be determined, given that Ras2 and PKA are also recruited to mitochondria by Hsp60 affecting their turnover (Portela and Rossi, 2020).

## 4. Material and methods

### 4.1. Yeast strains and growth conditions

Yeast strains used in this work are listed in Table 2 and were derived from HD56-5A, one of the parental strains of the common CEN.PK series (Arvanitidis and Heinisch, 1994), or its isogenic diploid DHD5 (Kirchrath et al., 2000). For cloning purposes and plasmid amplification, *E. coli* strain DH5α was used (Invitrogen, Karlsruhe, Germany). Standard procedures were followed for genetic manipulations of yeast and plasmid constructions (Rose et al., 1990). Complete sequences of all plasmids, modified chromosomal loci, and oligonucleotides employed are available upon request.

Rich medium (YEPD) was based on yeast extract (1% w/v) and peptone (2 % w/v), supplemented with 2% glucose (w/v). Synthetic media (SC) contained 0.67 % yeast nitrogen base (w/v) supplemented with ammonium sulfate, amino acids and bases as required (Rose et al., 1990), with 2% glucose (w/v, SCD) as a carbon source. Histidine concentration was raised from 2 to 4 mg/L, if necessary, to record growth curves. *E. coli* cells were grown in LB medium (yeast extract at 0.5% w/v, tryptone at 1% w/v, and sodium chloride at 1 % w/v), with the addition of 50 mg/L ampicillin or 25 mg/L kanamycin as required for plasmid selection.

### 4.2. Genetic manipulations

Deletion mutants were obtained by one-step gene replacements, using PCR products obtained with primers generating 40-50 bp of homology flanking the genomic target sequences, with selection for genetic markers as described (Gueldener et al., 2002). For complementation of auxotrophic markers with *KlURA3* (pJJH1286) or *KlLEU2* (pJJH1287) from *Kluyveromyces lactis,* modified plasmids were used, which carried the marker genes flanked by the *TEF2* promoter and the *TEF2* terminator from *Ashbya gossypii* and by two *loxP* sites. Alleles encoding hyper-active variants of the GTPases (*RHO5^G12V^* or *RAS2^G19V^*) were inserted at the native genetic loci by substitution of the respective deletion markers, using *SkHIS3* inserted into the respective 3’ non-coding regions as selection marker.

### 4.3. Determination of respiratory capacity

A Seahorse analyzer (Agilent Technologies Deutschland GmbH, Waldbronn, Germany) was employed to determine the respiratory capacity of the different yeast strains with the “Seahorse XF Cell Mito Stress Test” with a modification of the test employed for mammalian cells (Morris et al., 2023). It allows the measurement of the oxygen consumption rate in live cells. For the assay of yeast cells, cultures were grown overnight in SCD at 30°C with shaking (180 rpm). After dilution to an OD_600_ of 0.4 they were again incubated to reach an OD_600_ of 0.8. Three biological replicates and at least two technical replicates were determined, with the exception of a *rho5* deletion, where only one technical replicate was obtained for one of the three biological replicates.

### 4.4. Epistasis analyses

Strains with different deletion or mutant alleles were crossed by standard yeast genetic techniques (Rose et al., 1990), sporulated on plates with 1% potassium acetate, and subjected to tetrad analyses on YEPD plates using a Singer MSM400 micromanipulator (Singer Instruments, Somerset, UK). Plates were incubated for 3-5 days at 30°C as required for the specific mutant combinations, and scanned for documentation. The images were adjusted for brightness and contrast using ImageJ with the same settings for the entire plate, and colony sizes were determined with the analyze particles function of the program. At least 40 tetrads were separated for each cross and used to compare colony sizes after assigning the genotypes from marker analyses. The averaged sizes of wild-type segregants from each cross were set to 100% and the relative sizes of mutant segregants were averaged and calculated. Statistical analyses were carried out using the T.TEST function of Excel.

Growth curves of yeast strains under standard and oxidative stress conditions were recorded with a Varioscan Lux plate reader (ThermoFisher Scientific) as detailed in (Bischof et al., 2024b).

### 4.5. High-throughput analyses

RNA preparation, RNAseq and bioinformatic analyses were performed by StarSeq (Mainz, Germany). For this purpose, cells were grown in 50 mL SCD in the absence or presence of 0.8 mM hydrogen peroxide, inoculated from fresh overnight cultures to an OD_600_ of 0.2 and grown for another two generations to an OD_600_ of 0.8 at 30°C with shaking at 180 rpm. Cells were collected by centrifugation, frozen in liquid nitrogen and shipped on dry ice.

Proteomes were obtained from mass spectrometry analyses. Therefore, yeast cells were grown in 25 mL SCD, which were inoculated from a fresh overnight culture to an OD_600_ of 0.2 and allowed to grow for another two generations at 30°C and shaking at 180 rpm. Samples were extracted using the iST kit according to the instructions of the manufacturer (Preomics GmbH, Martinsried, Germany). Mass spectrometry using label-free quantification (LFQ) was performed at the Mass Spectrometry Equipment Center of the Department of Biology/Chemistry at the "CellNanOS" research center of the University of Osnabrück. For this purpose, dried peptides were resuspended in 10 µL LC-Load buffer and 2 µL were used to perform reversed-phase chromatography on a Thermo Ultimate 3000 RSLCnano system connected to a TimsTOF HT mass spectrometer (Bruker Corporation, Bremen) through a Captive Spray Ion source. Peptides were separated on an Aurora Gen3 C18 column (25 cm x 75 µm x 1.6 µm) with CSI emitter (Ionoptics, Australia) at temperature of 40°C. Peptides from the column were eluted via a linear gradient of acetonitrile from 10-35% in 0.1% formic acid (v/v) for 44 min at a constant flow rate of 300 nL/min following a 7 min increase to 50%, and finally, 4 min to reach 85% buffer B. Eluted peptides were then directly electro sprayed into the mass spectrometer at an electrospray voltage of 1.5 kV and 3 L/min dry gas.

The MS settings of the timsTOF were adjusted to positive Ion polarity with a MS range from 100 to 1700 m/z. The scan mode was set to PASEF. The ion mobility was ramped from 0.7 Vs/cm^2^ to 1.5 in 100 ms. The accumulation time was also set to 100 ms. 10 PASEF ramps per cycle resulted in a duty cycle time of 1.17 s. The target intensity was adjusted to 14,000, the intensity threshold to 1,200. The dynamic exclusion time was set to 0.4 min to avoid repeated scanning of the precursor ions, their charge state was limited from 0 to 5. The resulting data were analysed with PeaksOnline (BSI, Canada) version 11, employing the corresponding Yeast FASTA databases. Precursors were ranging from 600 to 6,000 Da. As modifications carbamidomethylation (C) and oxidation (M) were chosen. DDA-MBR were performed with MS tolerance of 10 ppm and IM tolerance of 0.05 (1/k0).

The obtained data from RNA-sequencing and mass spectrometry were processed using the standard Excel program. Cut-offs were applied at p-values less than 0.05 and at least a twofold change in expression. Proteins detected by mass spectrometry were only considered, if at least one unique peptide was repeatedly found. It should be noted that while some proteins could not be distinguished in the mass spectrometry data due to their high amino acid similarities (e.g. some members of the hexose transporter family), their encoding RNAs were clearly distinguished by RNAseq.

## Acknowledgements

We thank Rosaura Rodicio for critical reading of the manuscript and sharing her expertise in the regulation of yeast carbohydrate metabolism.

