## Supplementary figures 1-3 for "The Small GTPase Rho5 – Yet Another Player in Yeast Glucose Signaling"

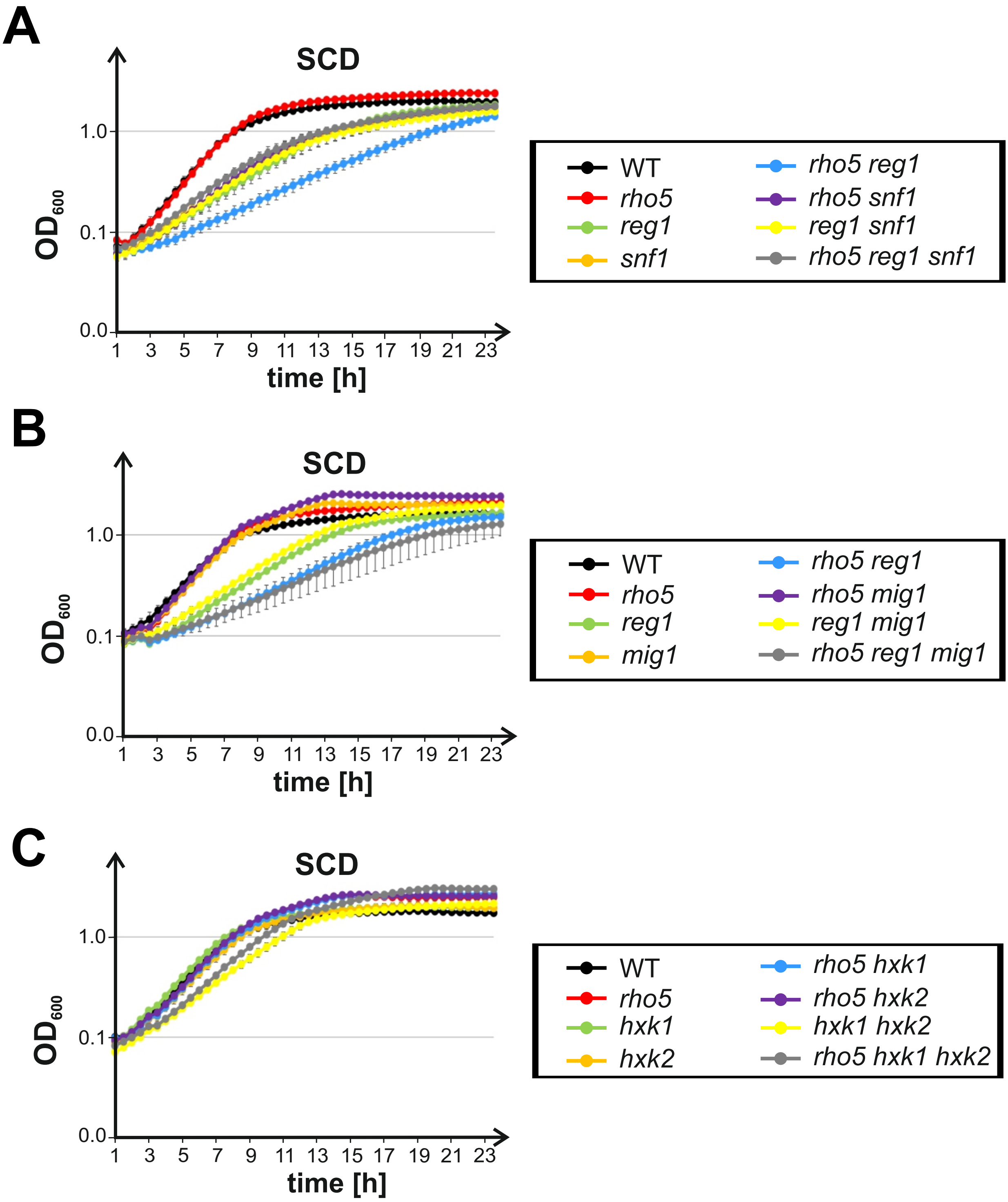


**Figure S1.** Epistasis analyses based on the growth of strains carrying mutations in genes encoding components of the SNF1 signalling pathway in combination with *RHO5* variants. Growth curves were recorded on synthetic medium with 2% glucose (SCD; supplemented with 4 mg/L histidine for strains with a histidine auxotrophy) as indicated. Error bars give the standard deviations at each time point obtained from at least two biological and two technical replicates from parallel measurements for each curve (i.e. two independent isogenic segregants were measured, with two independent inoculates, each). Genotypes of the otherwise isogenic strains are listed in Table 2 (main text). Strains employed were A) wild type (FSO71-1A and FSO71-7B) *rho5* (FSO71-2A and FSO71-15A) *reg1* (FSO71-10B and FSO71-1D) *snf1* (FSO71-5D and FSO71-9C) *rho5 reg1* (FSO71-9D and FSO71-15D) *rho5 snf1* (FSO71-1B and FSO71-6B) *reg1 snf1* (FSO71-15B and FSO71-2C) *rho5 reg1 snf1* (FSO71-7C and FSO71-9B). B) wild type (FSO86-7A and FSO86-2A) *rho5* (FSO75-4D and FSO75-7D) *reg1* (FSO71-10B and FSO71-1D) *mig1* (FSO90-7A and FSO90-2A) *rho5 reg1* (FSO71-9D and FSO71-15D) *rho5 mig1* (FSO90-1A and FSO90-8B) *reg1 mig1* (FSO79-4C and FSO79-8C) *rho5 reg1 mig1* (FSO90-3D and FSO90-6D). C) wild type (FSO55-9A and FSO55-9B) *rho5* (FSO56-1A and FSO56-2A) *hxk1* (FSO56-3A and FSO56-1B) *hxk2* (FSO56-1C and FSO56-3C) *rho5 hxk1* (FSO56-2B and FSO56-4A) *rho5 hxk2* (FSO56-4B and FSO56-9C) *hxk1 hxk2* (FSO56-8D and FSO56-4D) *rho5 hxk1 hxk2* (FSO56-6D and FSO56-1D).

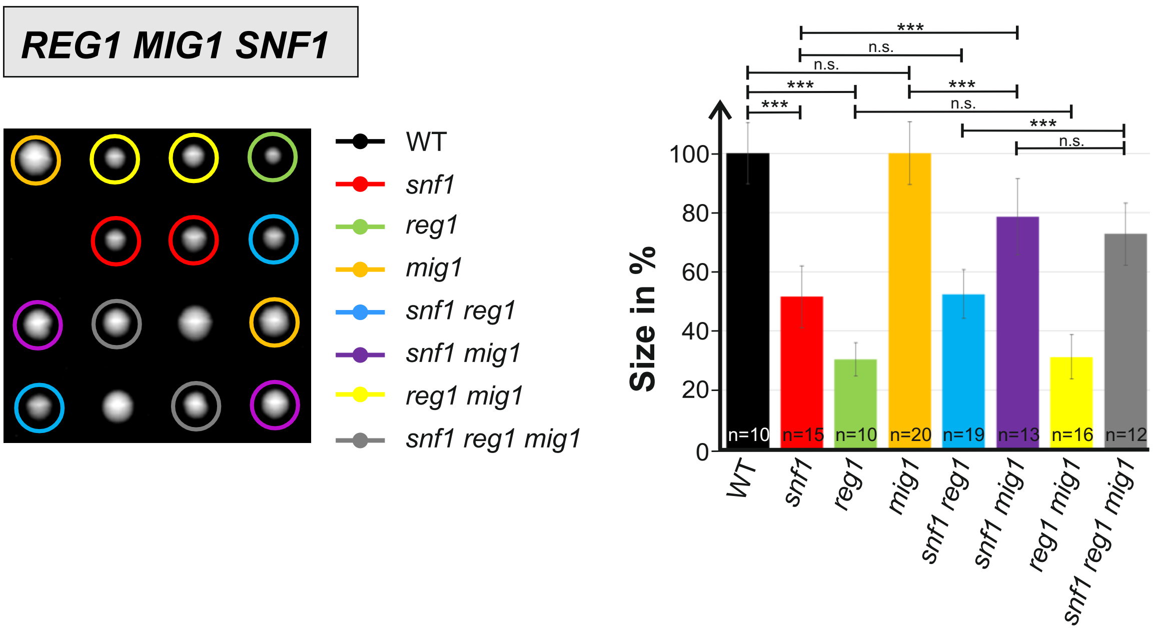


**Figure S2.** Epistasis analyses based on growth of segregants from tetrad analyses on rich medium (YEPD). Plates four exemplary tetrads are shown, with colored circles designating different combinations of gene deletions as indicated. Colony sizes for each combination (determined from pixel area and given as percentage from wild type set at 100%) were determined from 29 tetrads and quantified in the columns of the diagram at the right (n = total number of segregants obtained for each genotype. Error bars are indicated for each mutant combination. Three asterisks indicate highly significant differences with p-values below 0.001; n.s. = not significant). Diploids analyzed were from the cross of a strain carrying a *reg1 mig1* double deletion (FSO79-8C) with one carrying a *snf1* deletion (HOD201-2D).

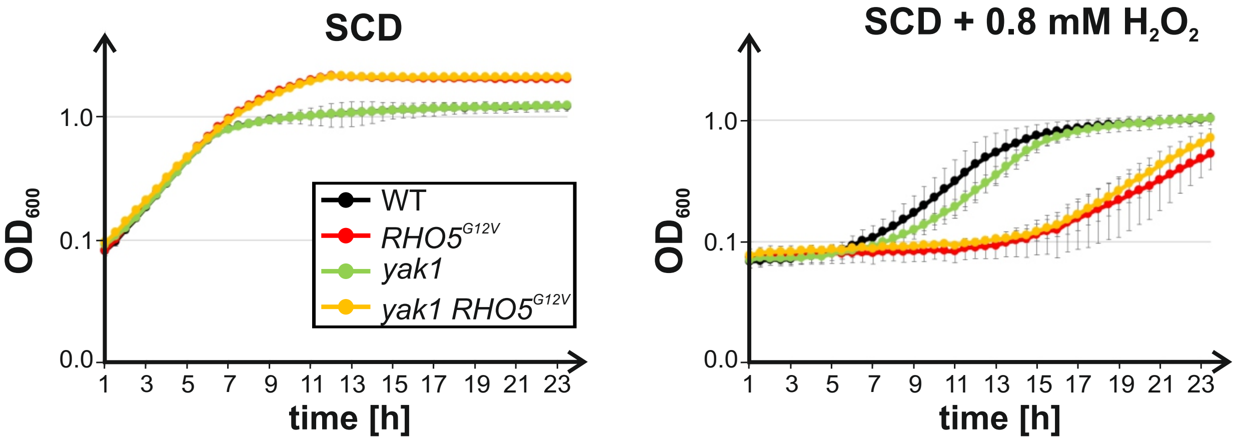


**Figure S3.** Epistasis analyses of *rho5* and *yak1* mutant combinations based on their sensitivity towards hydrogen peroxide. Growth curves were recorded on synthetic medium with 2% glucose (SCD), as described in materials and methods of the main text, with or without hydrogen peroxide as indicated. Errot bars give the standard deviations at each time point obtained from at least two biological and two technical replicates. Genotypes of the otherwise isogenic strains are listed in Table 2. Strains employed were wild type (FSO35-2A and FSO35-4A), *RHO5*$^{G12V}$, (FSO88-3A and FSO88-2B), *yak1* (FSO88-2C and FSO88-6A), *RHO5*$^{G12V}$ *yak1* (FSO88-2A and FSO88-1C).
